# Cholinergic activation of HLH-30/TFEB promotes *C. elegans* infection resilience

**DOI:** 10.64898/2026.09.18.752639

**Authors:** Swarupa Mallick, Xavier Gonzalez, Khursheed A. Wani, Sid A. Labed, Javier E. Irazoqui

**Affiliations:** Department of Microbiology, University of Massachusetts Chan Medical School, Worcester, MA 01605, United States of America

**Keywords:** muscarinic acetylcholine receptor, *Staphylococcus aureus*, innate host defense, neuroimmune signaling, Gαq signaling, protein kinase D

## Abstract

Innate host defense depends on transcriptional programs, which support survival through two strategies that can vary independently. Resistance lowers pathogen burden, while resilience limits the damage a given burden causes. In *Caenorhabditis elegans* infected with *Staphylococcus aureus*, the highly conserved transcription factor HLH-30/TFEB drives most of the host response, which mediates infection resilience. We previously showed that EGL-30/Gαq, PLC-1/PLCε, and DKF-1/PRKD activate HLH-30/TFEB during infection. The same phospholipase C and protein kinase D step activates TFEB in mouse macrophages infected with *Salmonella* or *S. aureus*. However, the input that drives the EGL-30/Gαq, PLC-1/PLCε, and DKF-1/PRKD module to activate HLH-30/TFEB during infection remained unknown. Here we report that acetylcholine, acting on muscarinic receptors, is one such input. Atropine and scopolamine, two muscarinic antagonists in clinical use, as well as muscarinic receptor gene silencing blunted HLH-30/TFEB activation by infection. Moreover, a mutation of *unc-17/SLC18A3* that lowers acetylcholine release from neurons impaired HLH-30/TFEB activation and induction of its target genes, while pathogen burden remained high. In uninfected animals, muscarinic agonist arecoline was sufficient to activate HLH-30/TFEB, in a muscarinic receptor-dependent manner. Nicotine, in contrast, had no effect. The agonist also lengthened survival of infection without lowering pathogen burden, in an HLH-30/TFEB-dependent manner. Silencing *egl-30/GNAQ*, *plc-1/PLCE1*, or *dkf-1/PRKD1-3* cut activation by the agonist just as it cut activation by infection, showing that both stimuli require the same module. In mammals the link documented so far between acetylcholine and TFEB runs through nicotinic receptors, so whether the muscarinic route is conserved remains open. Thus, we favor a model in which infection drives neurons to release acetylcholine onto the intestinal epithelium, where muscarinic receptors acting through EGL-30/Gαq, PLC-1/PLCε, and DKF-1/PRKD activate HLH-30/TFEB and strengthen infection resilience.

**Author Summary:** Animals survive an infection in two ways, often at once. They can kill the microbe, and they can limit the harm it does while it is still there. We study the second mechanism in the roundworm *Caenorhabditis elegans*, which fights bacteria with many of the same genes people do. A protein called HLH-30, known as TFEB in humans, switches on most of the genes this animal needs to withstand infection by *Staphylococcus aureus*. Which signals switch HLH-30 on during an infection has been unclear. We found that one of the signals comes from the nervous system. During infection, neurons release acetylcholine, a common neurotransmitter. Acetylcholine reaches the gut lining, where receptors pass the message inward and switch HLH-30 on. Drugs used in the clinic to block those receptors keep HLH-30 off, and so does removing the receptors themselves. A drug that stimulates the same receptors switches HLH-30 on even with no infection, and it helps animals survive infection without killing any more bacteria. The same receptors and the same protein are present in people. Whether this signal can be tuned to help a patient tolerate an infection is an important question to research in the future.

## Introduction

Innate host defense is an evolutionarily ancient feature of animal life, conserved across animals as distant as nematodes and mammals. The nematode *Caenorhabditis elegans* is a powerful system for studying this defense, because human bacterial pathogens kill nematodes through virulence factors that they also use in mammals, and because the animal defends itself through signaling pathways, such as the p38 mitogen-activated protein kinase (MAPK) cassette, that are conserved in mammals [1–3]. Much of this defense is transcriptional, because infection drives large changes in gene expression, and these transcriptional programs are central to host defense [4]. These transcriptional programs support host survival through two distinct strategies, resistance and resilience. Resistance lowers pathogen burden. Resilience limits the damage that a given burden causes, without clearing the pathogen [5, 6]. Resilience is frequently termed tolerance, a usage we avoid here to prevent confusion with immunological tolerance, which denotes acquired unresponsiveness to an antigen [7]. Resistance and resilience are separable, so the two can vary independently [5, 8]. Because the transcriptional host response drives both [4, 9], the transcription factors that control it shape the outcome of infection, and with it the course of infectious and inflammatory disease [9, 10].

HLH-30 is one such transcription factor, the *C. elegans* ortholog of mammalian TFEB, a basic helix-loop-helix leucine-zipper transcription factor conserved from nematodes to mammals [9, 11]. TFEB is a master regulator of the autophagy and lysosomal gene network [12, 13]. In *C. elegans*, HLH-30/TFEB becomes active soon after infection with human pathogenic bacterium *Staphylococcus aureus* and drives nearly 80% of the host response, including the antimicrobial and autophagy genes the animal needs for resilience to infection, and TFEB is likewise activated in infected mouse macrophages [9]. This immune role is broadly conserved, because TFEB and its paralog TFE3 also drive proinflammatory programs in mammalian macrophages [14–16], and TFEB is required for the enhanced bacterial killing that follows Fcγ receptor engagement [17]. The activities of HLH-30 and TFEB depend on their location in the cell. Each factor stays in the cytosol when it is inactive, and it moves into the nucleus to switch on its target genes [11, 12, 18, 19]. In mammals, this nuclear entry is known to be set by nutrient and mTORC1 signaling [18, 19]. However, the signals that activate HLH-30/TFEB during infection remain poorly understood [20].

We previously began to genetically define how infection activates HLH-30/TFEB. In *C. elegans* infected with *S. aureus*, EGL-30/Gαq acts through PLC-1/PLCε and DKF-1/PRKD to activate HLH-30/TFEB [20]. The same phospholipase C and protein kinase D cassette activates TFEB in *Salmonella*- and *S. aureus-*infected mouse macrophages, so it has been conserved across roughly one billion years of evolution [20]. Yet what activates this cassette during infection remains unknown, and the receptor that acts upstream of EGL-30/Gαq has not been identified [20].

Acetylcholine is a conserved neurotransmitter that signals through two receptor classes with distinct mechanisms. Muscarinic receptors are metabotropic and couple to Gαq/11 [21], whereas nicotinic receptors are ionotropic ligand-gated ion channels [22]. EGL-30 is a Gαq protein, so muscarinic receptors couple to the same class of G protein that initiates the cassette upstream of HLH-30/TFEB. Whether muscarinic signaling engages that cassette is unknown. *C. elegans* expresses three muscarinic-type G-protein-coupled acetylcholine receptors, GAR-1/mAChR, GAR-2/mAChR, and GAR-3/mAChR [23, 24], and in its nervous system it expresses UNC-17/VAChT, the vesicular acetylcholine transporter that loads acetylcholine into synaptic vesicles for release in the nervous system [25].

We recently connected cholinergic signaling to host defense in *C. elegans* [10]. Infection triggers acetylcholine release from neurons. That acetylcholine stimulates muscarinic signaling in the intestinal epithelium, acting through GAR-2/mAChR and GAR-3/mAChR. Through EGL-30/Gαq, those receptors induce Wnt expression, which drives antimicrobial gene expression, thus protecting the animal [10]. With this precedent in mind, we hypothesized that acetylcholine might also regulate HLH-30/TFEB. Here we report that cholinergic stimulation is sufficient to drive HLH-30/TFEB into the nucleus and to enhance infection resilience in an *hlh-30*/TFEB-dependent manner. We further show that endogenous acetylcholine acting through muscarinic receptors is necessary for HLH-30/TFEB activation during infection, and that this control runs through the EGL-30/Gαq, PLC-1/PLCε, and DKF-1/PRKD pathway. Thus, neural cholinergic control of the conserved transcription factor TFEB strengthens host resilience to infection.

## Results

### Muscarinic acetylcholine receptors are required for HLH-30/TFEB activation during infection

Infection with *S. aureus* drives HLH-30/TFEB into the nucleus within thirty minutes, and the transcriptional program that follows is necessary for the animal to survive infection [9]. To determine whether muscarinic receptor function is required for HLH-30/TFEB activation, we treated animals expressing GFP-tagged HLH-30 (HLH-30::GFP) with receptor antagonists atropine and scopolamine before infection. Animals were pretreated on solid medium for sixteen hours, then infected for thirty minutes (**Figure 1A**). Because the intestine is the infected organ in the *S. aureus* infection, we counted GFP+ nuclei within the posterior intestine (**Figure 1B**), which is amenable to live light microscopy and where HLH-30/TFEB activation is most robust during infection. In vehicle controls infected with *S. aureus*, nuclear HLH-30::GFP was observed in various tissues, including neurons, epidermis (hypodermis), body wall muscle, and intestinal epithelium (**Figure 1C**), consistent with our previous findings.

**Figure 1.**
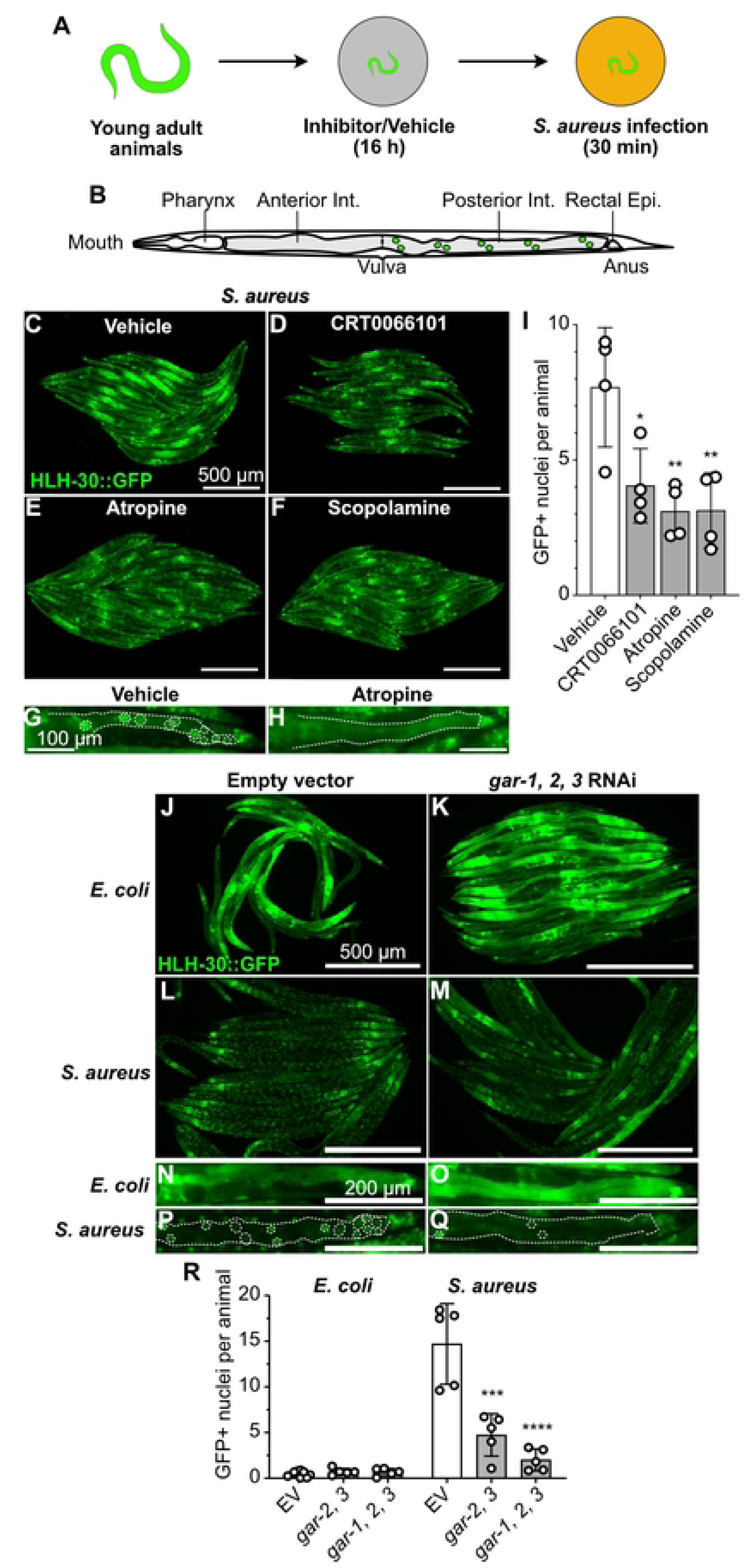
Muscarinic receptors are required for HLH-30/TFEB nuclear localization during infection. (A) Experimental scheme. Young adult animals were incubated with inhibitor or vehicle for 16 h and subsequently infected with *S. aureus* SH1000 for 30 min. (B) Schematic of an adult animal with anatomical landmarks, the pharynx, the anterior intestine, the vulva, the posterior intestine, and the rectal epithelium. Green circles mark GFP-positive nuclei, which are drawn posterior to the vulva only. (C–F) Representative epifluorescence micrographs of HLH-30::GFP animals infected with *S. aureus* SH1000 after treatment with vehicle (C), 5 μM CRT0066101 (D), 5 mM atropine (E), or 5 mM scopolamine (F). Scale bars, 500 μm. (G, H) Higher magnification of the posterior intestine of animals treated with vehicle (G) or atropine (H). Dotted lines outline the intestine, and dotted circles in (G) exemplify GFP-positive nuclei identified in FIJI by thresholding and particle detection. Some visible nuclei touch, or do not reach the conservative threshold and therefore the intensity falls below the positivity cutoff. Scale bars, 100 μm. (I) Quantitative analysis of (C–F). Data are mean ± SD (4 independent biological replicates, N = 20 to 40 animals per condition). * *P* ≤ 0.05, ** *P* ≤ 0.01 versus vehicle (one-way ANOVA followed by Dunnett’s multiple comparisons test). (J–M) Representative epifluorescence micrographs of HLH-30::GFP animals reared on *E. coli* HT115(DE3) carrying empty vector (J, L) or *gar-1*/*CHRM1-5*, *gar-2*/*CHRM1-5*, and *gar-3*/*CHRM1-5* triple RNAi (K, M), and subsequently fed *E. coli* OP50 (J, K) or infected with *S. aureus* SH1000 (L, M) for 30 min at 25 °C. Scale bars, 500 μm. (N–Q) Higher magnification of the intestine of animals treated as in (J–M), fed *E. coli* OP50 (N, O) or infected with *S. aureus* SH1000 (P, Q). In (P) and (Q), dotted lines outline the intestine and dotted circles exemplify GFP-positive nuclei. Scale bars, 200 μm. (R) Quantitative analysis of HLH-30::GFP nuclear localization in animals reared on *E. coli* HT115(DE3) carrying empty vector, *gar-2*/*CHRM1-5* and *gar-3*/*CHRM1-5* double RNAi, or *gar-1*/*CHRM1-5*, *gar-2*/*CHRM1-5*, and *gar-3*/*CHRM1-5* triple RNAi, and subsequently fed *E. coli* OP50 or infected with *S. aureus* SH1000. Data are mean ± SD (5 independent biological replicates, N = 20 to 40 animals per condition). ns, not significant. *** *P* ≤ 0.001, **** *P* ≤ 0.0001 versus infected empty vector (one-way ANOVA followed by Dunnett’s multiple comparisons test).

The antagonists had mixed effects on HLH-30/TFEB localization, but showed a greater effect than the protein kinase D inhibitor CRT0066101 used as positive control (**Figure 1D**) [20]. Nuclear HLH-30::GFP was variably discernable in antagonist-treated animals, mostly in the extra-intestinal tissues (**Figure 1D to 1F**), whereas the intestinal epithelium showed a noticeable and partially penetrant reduction in GFP+ nuclei (**Figure 1G and 1H**). CRT0066101 lowered nuclear HLH-30/TFEB by about 45% relative to a vehicle mean of ∼8 GFP+ nuclei per animal (**Figure 1D and 1I**; adjusted *P* = 0.0162). By comparison, atropine and scopolamine demonstrated a slightly stronger effect, reducing nuclear HLH-30/TFEB by about 60% (**Figure 1C**, **1E**, **1F, and 1I**; adjusted *P* = 0.0035 and *P* = 0.0036). These results showed that two different muscarinic antagonists in clinical use impair HLH-30 nuclear localization during infection.

To further test the role of muscarinic signaling, we silenced the three *C. elegans* muscarinic acetylcholine receptor genes, *gar-1/CHRM1-5*, *gar-2/CHRM1-5*, and *gar-3/CHRM1-5*, in HLH-30::GFP animals. We counted GFP+ nuclei in the posterior intestinal epithelium after thirty minutes of infection (**Figure 1J to 1R**). Animals fed nonpathogenic *Escherichia coli* showed almost no nuclear HLH-30/TFEB, whatever the RNAi treatment (**Figure 1J, K, N, O, and R**). In contrast, infection raised the count to ∼15 nuclei per animal in empty vector controls (**Figure 1L**, **1P, and 1R**). Silencing *gar-2/CHRM1-5* and *gar-3/CHRM1-5* together lowered it by about 70% (**Figure 1R**; adjusted *P* = 0.0003), while silencing all three receptors lowered it by about 85% (**Figure 1M**, **1Q, and 1R**; adjusted *P* < 0.0001). As with the antagonists, the RNAi silencing disrupted HLH-30::GFP nuclear localization mostly in the intestinal epithelium and hypodermis, while nuclear signal was still observable in muscle and neurons. Thus, while not affecting its nuclear localization on nonpathogenic food, muscarinic receptors are genetically required for HLH-30/TFEB activation by infection in the intestinal epithelium. Jointly, these results suggest that pharmacological or genetic inhibition of the muscarinic receptors impairs HLH-30 activation by infection.

### Endogenous acetylcholine release is required for HLH-30/TFEB activation during infection

To determine whether endogenous acetylcholine release is required for HLH-30 activation by infection, we examined animals carrying a hypomorphic loss of function mutation in *unc-17/SLC18A3*, which encodes the vesicular acetylcholine transporter (VAChT). UNC-17/VAChT loads acetylcholine into synaptic vesicles, and animals lacking *unc-17/SLC18A3* function release less of it [10]. While on nonpathogenic food they showed little HLH-30::GFP nuclear localization, after thirty minutes of infection, wild type animals showed clear nuclear localization throughout the body, as expected (**Figure 2A to 2D**). Noninfected *unc-17/SLC18A3* mutants were indistinguishable from wild type, but infection produced about 50% HLH-30::GFP nuclear localization in both the head and the posterior intestine compared to wild type (**Figure 2E to 2J**). These results show that endogenous acetylcholine release is required for full HLH-30::GFP nuclear localization throughout the body during infection.

**Figure 2.**
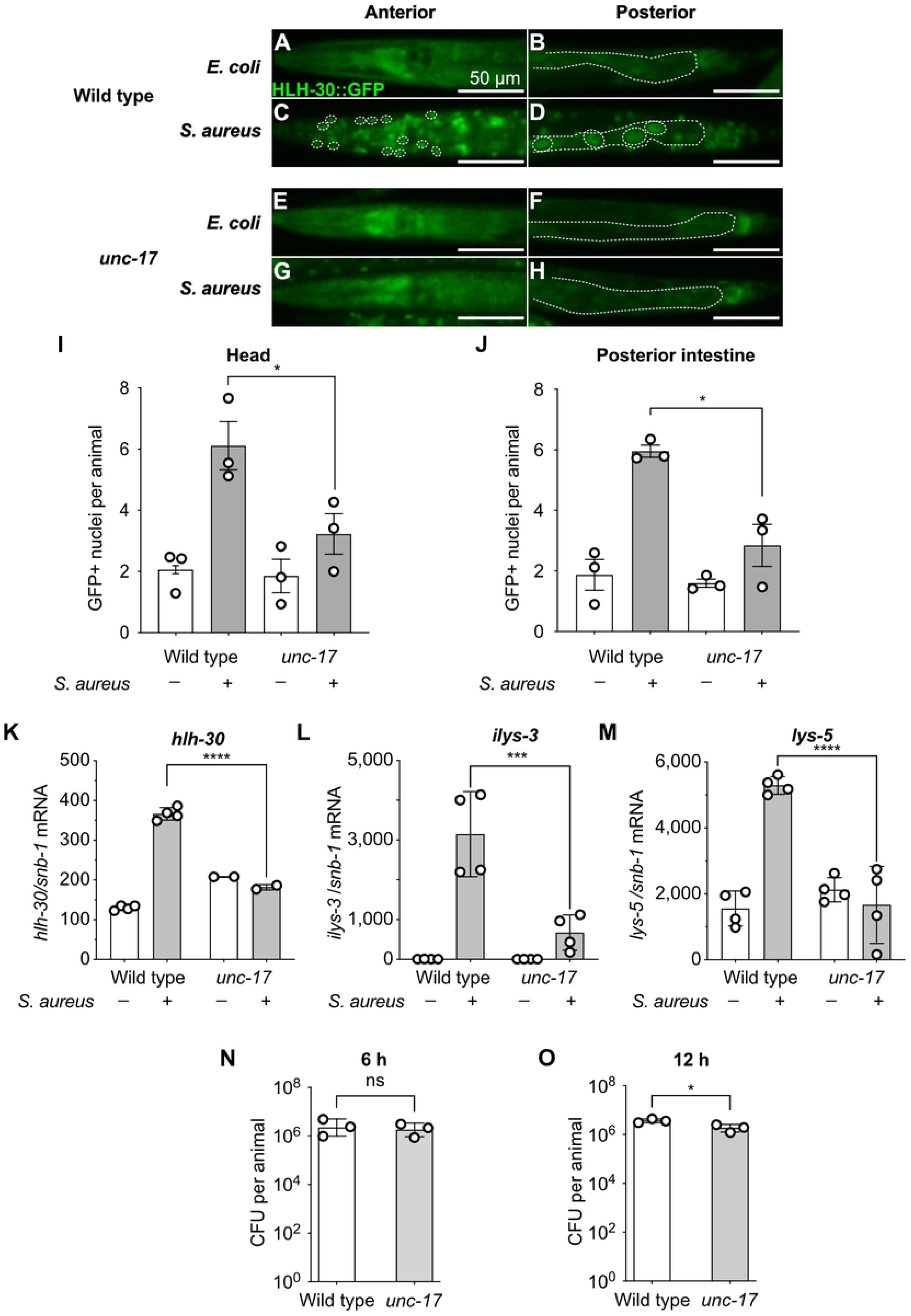
Endogenous acetylcholine is required for HLH-30/TFEB activation during infection. (A–D) Representative epifluorescence micrographs of the anterior (A, C) and posterior (B, D) of wild type animals expressing HLH-30::GFP, fed *E. coli* OP50 (A, B) or infected with *S. aureus* SH1000 (C, D) for 30 min at 25 °C. Dotted lines outline the intestine in (B) and (D), and dotted circles in (C) and (D) exemplify GFP-positive nuclei. Scale bars, 50 μm. (E–H) As in (A–D), for *unc-17(e245)*/*SLC18A3* deficient animals fed *E. coli* OP50 (E, F) or infected with *S. aureus* SH1000 (G, H) for 30 min at 25 °C. Dotted lines outline the intestine in (F) and (H). Scale bars, 50 μm. (I) Quantitative analysis of HLH-30::GFP nuclear localization in the head of wild type and *unc-17(e245)*/*SLC18A3* deficient animals fed *E. coli* OP50 or infected with *S. aureus* SH1000 for 30 min at 25 °C. Data are means ± SEM from three biological replicates, N = 20 to 40 animals per condition. * *P* ≤ 0.05 (unpaired two-tailed *t* test). (J) As in (I), scored in the posterior intestine. Data are means ± SEM from three biological replicates, N = 20 to 40 animals per condition. * *P* ≤ 0.05 (unpaired two-tailed *t* test). (K–M) RT-qPCR of *hlh-30*/*TFEB* (K), *ilys-3* (L), and *lys-5* (M) in wild type and *unc-17(e245)*/*SLC18A3* deficient animals fed *E. coli* OP50 or infected with *S. aureus* SH1000 for 8 h. Results are normalized to *snb-1*. Data are mean ± SD (*hlh-30*, 4 independent biological replicates for wild type and 2 for *unc-17(e245)*; *ilys-3* and *lys-5*, 4 independent biological replicates each). **** *P* ≤ 0.0001, *** *P* ≤ 0.001 (ordinary one-way ANOVA with Šídák multiple comparisons test). (N, O) Intestinal accumulation of *S. aureus* SH1000 in wild type and *unc-17(e245)*/*SLC18A3* deficient animals after 6 h (N) or 12 h (O) of infection at 25 °C, expressed as colony-forming units per animal on a logarithmic axis. Data are means ± SD of 3 independent biological replicates; each replicate is the mean of the log10 of its technical tubes, with ten animals pooled per tube (two or three tubes per genotype). ns, not significant in (N). In (O), * *P* ≤ 0.05 (paired two-tailed t test on the log10 values).

Nuclear localization is a proxy for HLH-30/TFEB activation. To determine whether cholinergic signaling controls HLH-30/TFEB activity, we measured transcript abundance in wild-type and *unc-17/SLC18A3* animals after eight hours of infection, the interval over which the host response develops [9]. Infection induced *hlh-30*/TFEB, which is subject to autoregulation [26], about 2.8-fold in wild-type animals, and that induction was lost in *unc-17/SLC18A3* mutants (**Figure 2K**; adjusted *P* < 0.0001 for the wild-type induction and adjusted *P* = 0.1438 for the mutant). Two HLH-30/TFEB-dependent lysozyme genes behaved the same way. Induction of *ilys-3* was reduced about five-fold, and induction of *lys-5* was abolished (**Figure 2L and 2M**; *ilys-3* adjusted *P* < 0.0001 in wild type and *P* = 0.4268 in the mutant, *lys-5* adjusted *P* < 0.0001 in wild type and *P* = 0.8311 in the mutant).

A smaller response might follow from a smaller stimulus, i.e. pathogen load. To exclude that possibility, we measured intestinal pathogen burden at two timepoints. At six hours, wild-type and *unc-17/SLC18A3* animals carried indistinguishable numbers of *S. aureus*, and by twelve hours *unc-17/SLC18A3* animals carried fewer, 1.8 × 10⁶ against 3.6 × 10⁶ colony-forming units per animal (**Figure 2N and 2O**; six hours *P* = 0.5551, twelve hours *P* = 0.026). Therefore, a smaller bacterial stimulus is unlikely to account for the observed transcriptional defect. Together, the data indicate that endogenous acetylcholine is required for HLH-30/TFEB activation in response to infection.

### Muscarinic agonists are sufficient to activate HLH-30/TFEB, and a nicotinic agonist is not

Selective drugs make it possible to distinguish the two acetylcholine receptor classes. Arecoline and oxotremorine act as muscarinic agonists in *C. elegans*, and both trigger GAR-2/mAChR and GAR-3/mAChR [10, 27]. Nicotine and carbachol both excite pharyngeal muscle through a nicotinic receptor [28], although carbachol is not selective, because it also stimulates GAR-3/mAChR in heterologous cells [29].

To determine whether cholinergic stimulation alone activates HLH-30/TFEB, we exposed noninfected animals to cholinergic agonists on solid medium for thirty minutes (**Figure 3A to 3E**). Arecoline raised nuclear HLH-30/TFEB about 13-fold over a vehicle mean of 0.42 nuclei per animal (**Figure 3B and 3F**; adjusted *P* < 0.0001). The broad-spectrum cholinergic agonist carbachol and the muscarinic-selective agonist oxotremorine produced smaller increases, of about six-fold, that were not distinguishable from one another (**Figure 3C**, **3D, and 3F**; adjusted *P* = 0.0221 and *P* = 0.0155). Nicotine, in contrast, did not activate HLH-30/TFEB (**Figure 3E and 3F**; adjusted *P* = 0.7348). Arecoline drove HLH-30/TFEB into nuclei in the head (around 2.5-fold) and in the posterior intestine (∼threefold, **Figure 3G to 3L**; head *P* = 0.0112, posterior intestine *P* = 0.0016) compared with vehicle. Arecoline is also sufficient to induce the HLH-30/TFEB antimicrobial genes *ilys-3* and *lys-5* in noninfected animals [10]. These data altogether show that acute muscarinic stimulation is sufficient to activate HLH-30/TFEB in the absence of a pathogen, on the timescale over which infection acts.

**Figure 3.**
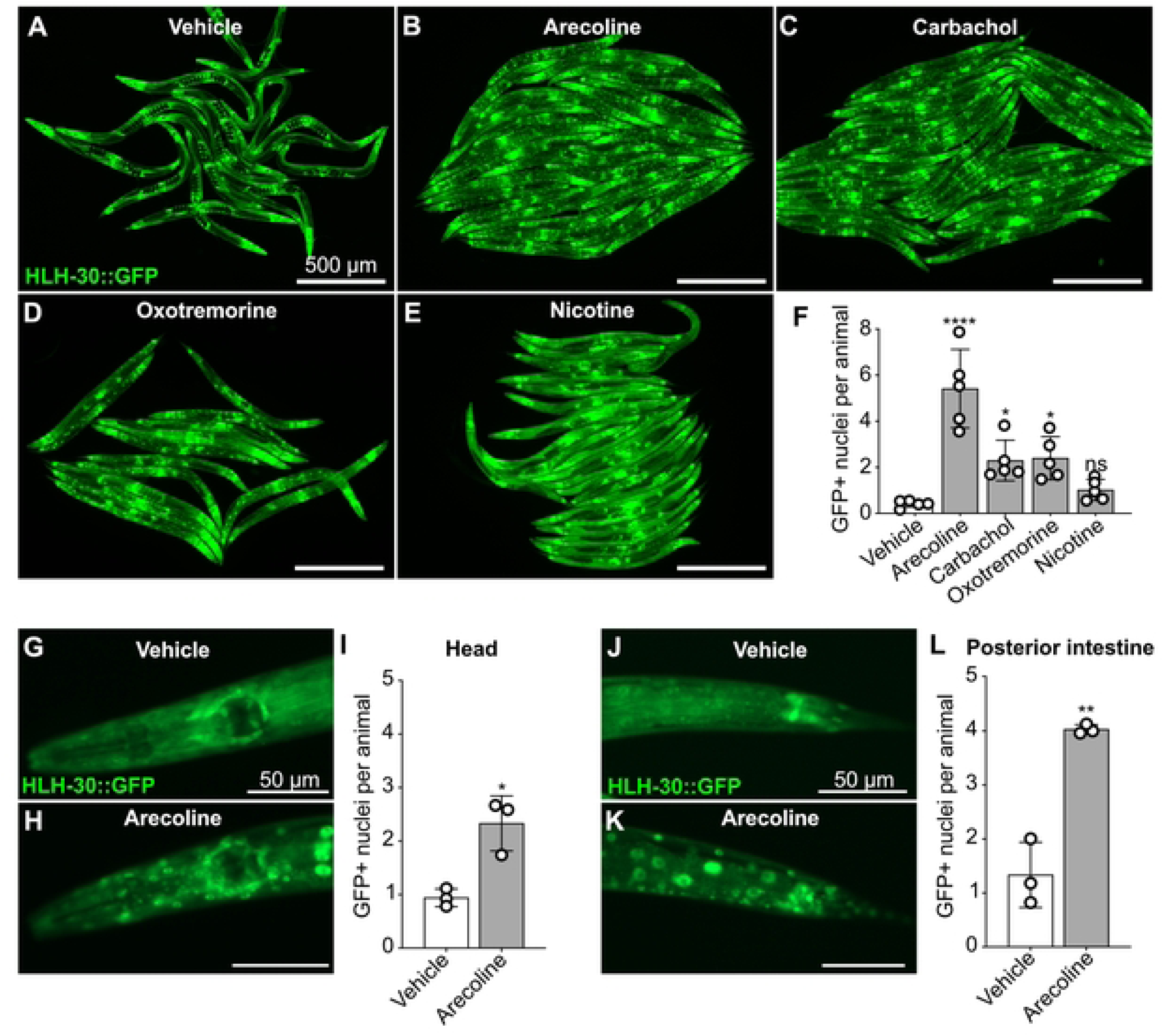
Cholinergic agonists drive HLH-30/TFEB into the nucleus. (A–E) Representative epifluorescence micrographs of HLH-30::GFP animals treated for 30 min at 25 °C on *E. coli* OP50, without infection, with vehicle (A), 10 mM arecoline (B), 10 mM carbachol (C), 10 mM oxotremorine (D), or 10 mM nicotine (E). Scale bars, 500 μm. (F) Quantitative analysis of (A–E). Data are mean ± SD (5 independent biological replicates, N = 20 to 40 animals per condition). ns, not significant. * *P* ≤ 0.05, **** *P* ≤ 0.0001 versus vehicle (one-way ANOVA followed by Dunnett’s multiple comparisons test). (G, H) Representative epifluorescence micrographs of the anterior end, or head, of HLH-30::GFP animals treated with vehicle (G) or 10 mM arecoline (H) for 30 min at 25 °C. Scale bars, 50 μm. (I) Quantitative analysis of HLH-30::GFP nuclear localization in the head of animals treated with vehicle or 10 mM arecoline for 30 min at 25 °C. Data are mean ± SD from three biological replicates, N = 23 to 52 animals per condition per condition. * *P* ≤ 0.05 (unpaired two-tailed *t* test). (J, K) Representative epifluorescence micrographs of the posterior end of HLH-30::GFP animals treated with vehicle (J) or 10 mM arecoline (K) for 30 min at 25 °C. Scale bars, 50 μm. (L) As in (I), scored in the posterior intestine. Data are mean ± SD from three biological replicates, N = 24 to 56 animals per condition. ** *P* ≤ 0.01 (unpaired two-tailed *t* test).

### Agonist-driven HLH-30/TFEB activation requires the same muscarinic receptors as infection

If arecoline works through muscarinic receptors, losing those receptors should abolish its effect. To test that prediction, we treated empty vector and *gar* RNAi animals with vehicle or the agonist (**Figure 4A to 4D**). Vehicle-treated animals showed little nuclear HLH-30/TFEB, whatever the RNAi treatment (**Figure 4A**, **4C, and 4E**), while arecoline raised the count in empty vector animals to about 10 nuclei per animal (**Figure 4B and 4E**). Silencing *gar-2/CHRM1-5* and *gar-3/CHRM1-5* lowered this by roughly 70%, and silencing all three receptors lowered it by about 85% (**Figure 4D and 4E**; adjusted *P* = 0.0063 and *P* = 0.0020). These results show that the muscarinic receptors are required for exogenous cholinergic activation of HLH-30/TFEB.

**Figure 4.**
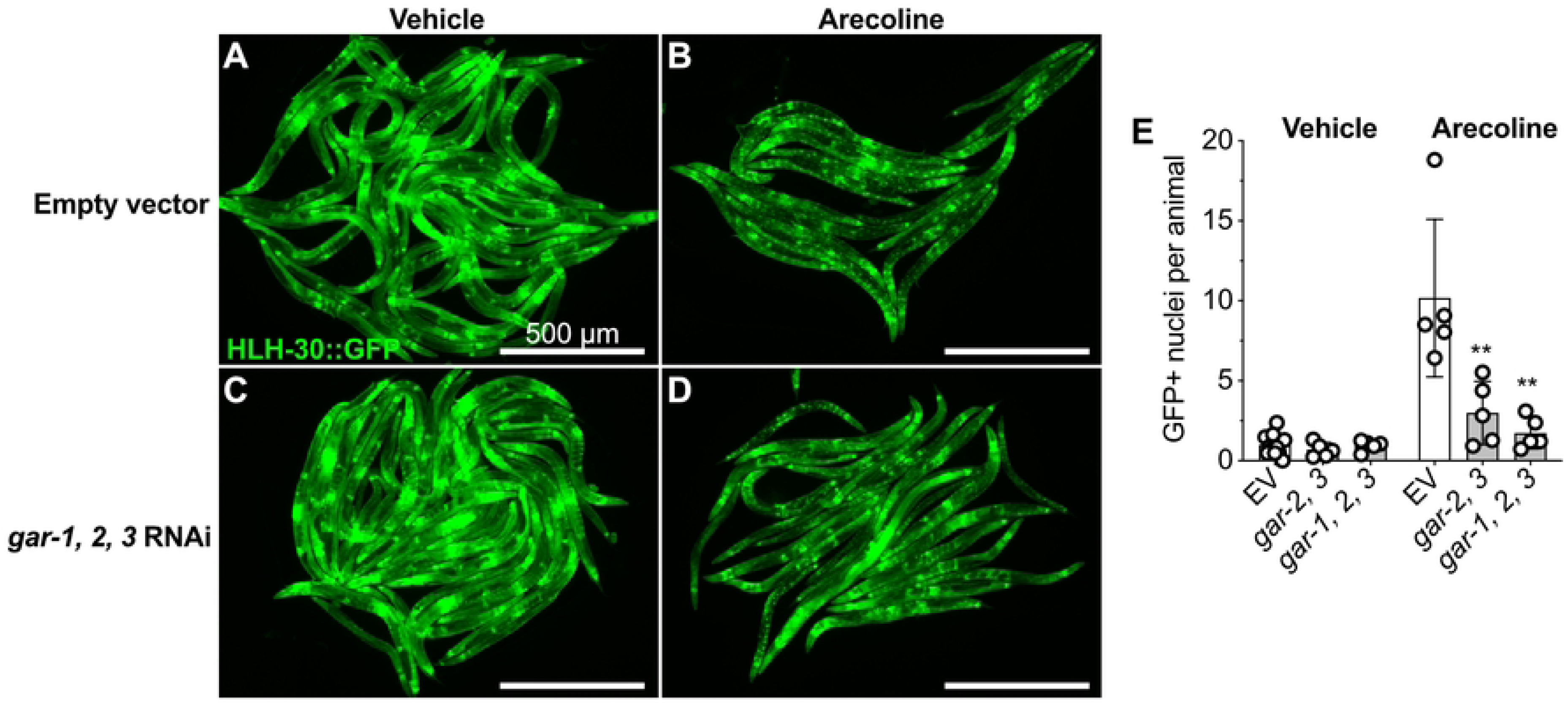
Muscarinic receptors are required for HLH-30/TFEB activation by arecoline. (A–D) Representative epifluorescence micrographs of HLH-30::GFP animals reared on *E. coli* HT115(DE3) carrying empty vector (A, B) or *gar-1*/*CHRM1-5*, *gar-2*/*CHRM1-5*, and *gar-3*/*CHRM1-5* triple RNAi (C, D), and subsequently treated with vehicle (A, C) or 10 mM arecoline (B, D) for 30 min at 25 °C. Scale bars, 500 μm. (E) Quantitative analysis of HLH-30::GFP nuclear localization in animals reared on *E. coli* HT115(DE3) carrying empty vector, *gar-2*/*CHRM1-5* and *gar-3*/*CHRM1-5* double RNAi, or *gar-1*/*CHRM1-5*, *gar-2*/*CHRM1-5*, and *gar-3*/*CHRM1-5* triple RNAi, and subsequently treated with vehicle or 10 mM arecoline. Data are mean ± SD (5 independent biological replicates, N = 20 to 40 animals per condition). ** *P* ≤ 0.01 versus arecoline-treated empty vector (one-way ANOVA followed by Dunnett’s multiple comparisons test).

### Muscarinic activation improves survival of infection and requires HLH-30/TFEB

To determine how muscarinic activation of HLH-30/TFEB affects the outcome of infection, we treated wild-type and *hlh-30*/*TFEB* mutant animals with arecoline and followed survival. Animals were treated for sixteen hours and then transferred to *S. aureus* (**Figure 5A**). Arecoline protected wild type animals compared to vehicle-treated controls (log-rank *P* < 0.0001), but such protection was negated by mutation of *hlh-30*/*TFEB* (**Figure 5B**). Arecoline did not change intestinal pathogen burden at twelve hours in either genotype (*P* = 0.9332 in wild type and *P* = 0.1238 in the mutant, **Figure 5C**). Therefore, arecoline improved survival without reducing the pathogen load.

**Figure 5.**
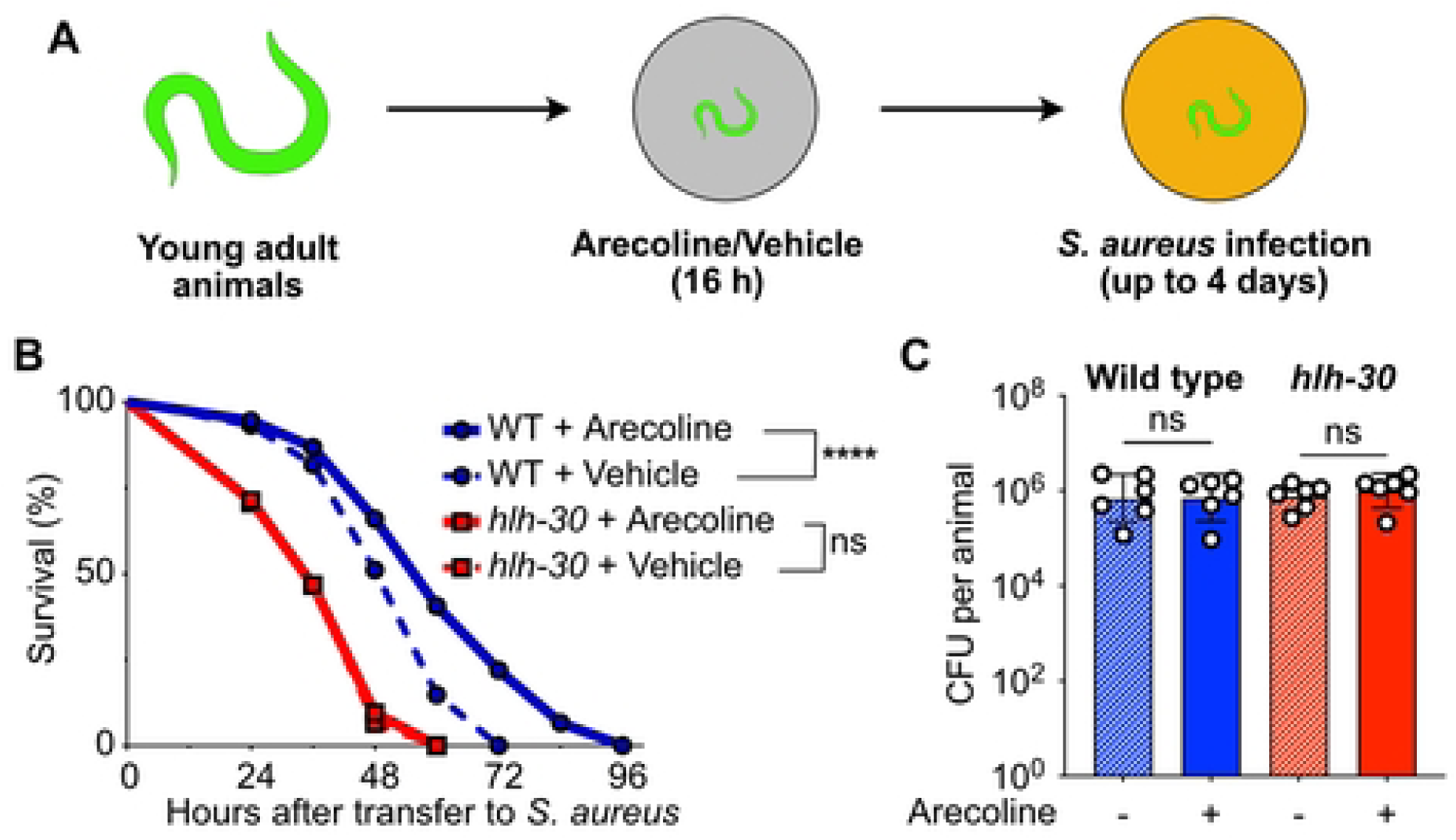
Cholinergic stimulation enhances survival of infection in an *hlh-30*/*TFEB*-dependent manner without lowering pathogen burden. (A) Experimental scheme. Young adult animals were incubated with arecoline or vehicle for 16 h and subsequently infected with *S. aureus* SH1000 for up to 4 days. (B) Survival of wild type and *hlh-30(tm1978)*/*TFEB* mutant animals treated as in (A) with 1 mM arecoline or vehicle. **** *P* ≤ 0.0001, n.s., not significant (Log-Rank test). Representative of four independent experiments. N = 88 to 94 animals per condition in the experiment shown. (C) Intestinal accumulation of *S. aureus* SH1000 in wild type and *hlh-30(tm1978)*/*TFEB* mutant animals treated with vehicle or 1 mM arecoline as in (A), after 12 h of infection, expressed as colony-forming units per animal on a logarithmic axis. Data are the mean and SD of 6 independent biological replicates; each replicate is the mean of the log10 of its technical tubes (two or three tubes per condition). ns, not significant (paired two-tailed *t* test on the log10 values, within genotype).

### HLH-30/TFEB activation by infection and by muscarinic agonist requires EGL-30/Gαq, PLC-1/PLCε, and DKF-1/PRKD

We previously showed that HLH-30/TFEB activation during infection requires EGL-30/Gαq, PLC-1/PLCε, and DKF-1/PRKD, acting downstream of an unidentified G protein-coupled receptor [20]. To verify the role of such cassette downstream of infection, we individually silenced *egl-30/GNAQ*, *plc-1/PLCE1*, and *dkf-1/PRKD1-3* in wild type animals prior to *S. aureus* infection. As expected, no RNAi treatment affected HLH-30/TFEB localization on nonpathogenic *E. coli*, while on *S. aureus* silencing *egl-30/GNAQ*, *plc-1/PLCE1*, and *dkf-1/PRKD1-3* reduced nuclear HLH-30/TFEB by about 50%, 60%, and 80% respectively (**Figure 6A**; adjusted *P* = 0.0296, *P* = 0.0052, and *P* = 0.0004).

**Figure 6.**
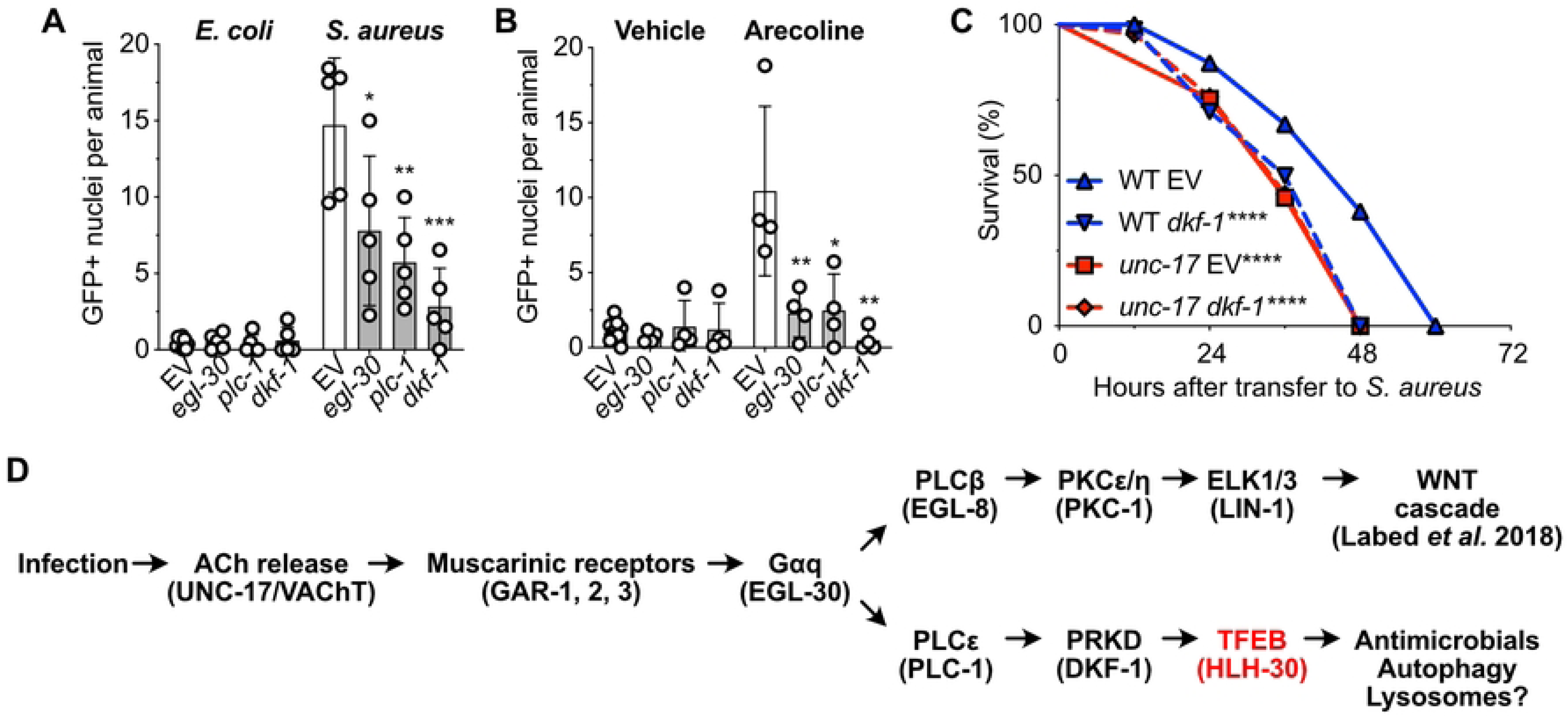
An EGL-30/Gαq, PLC-1/PLCε, and DKF-1/PRKD pathway links cholinergic signaling to HLH-30/TFEB. (A) Quantitative analysis of HLH-30::GFP nuclear localization in animals reared on *E. coli* HT115(DE3) carrying empty vector, *egl-30*/*GNAQ*, *plc-1*/*PLCE1*, or *dkf-1*/*PRKD* RNAi, and subsequently fed *E. coli* OP50 or infected with *S. aureus* SH1000 for 30 min. Data are mean ± SD (5 independent biological replicates, N = 20 to 40 animals per condition). * *P* ≤ 0.05, ** *P* ≤ 0.01, *** *P* ≤ 0.001 versus infected empty vector (one-way ANOVA followed by Dunnett’s multiple comparisons test). (B) As in (A), except that animals were treated with vehicle or 10 mM arecoline for 30 min at 25 °C. EV vehicle shows pooled results from 12 experiments performed at the same times as Fig. 4E. ** *P* ≤ 0.01, * *P* ≤ 0.05 versus arecoline-treated empty vector (one-way ANOVA followed by Dunnett’s multiple comparisons test). (C) Survival of wild type and *unc-17(e245)*/*SLC18A3* deficient animals reared on *E. coli* HT115(DE3) carrying empty vector or *dkf-1*/*PRKD* RNAi, and subsequently infected with *S. aureus* SH1000. **** *P* ≤ 0.0001 versus wild type empty vector (Log-Rank test). Representative of three independent trials. N = 61 to 96 animals per condition in the trial shown. (D) Working model. Infection triggers acetylcholine release, which requires UNC-17/VAChT. Acetylcholine acts on the muscarinic receptors GAR-1/mAChR, GAR-2/mAChR, and GAR-3/mAChR, which couple to EGL-30/Gαq. Signaling then branches. One branch runs through EGL-8/PLCβ, PKC-1/PKCε and PKCη, and LIN-1/ELK1 and ELK3 to the Wnt cascade [10]. The other branch runs through PLC-1/PLCε and DKF-1/PRKD to HLH-30/TFEB, which drives antimicrobial, autophagy, and possibly lysosomal programs.

If the muscarinic receptors function above this module, agonist-driven activation should require the same three components. To test that prediction, we treated the same RNAi animals with arecoline. Arecoline raised nuclear HLH-30/TFEB to about ten nuclei per animal in empty vector animals, and silencing *egl-30/GNAQ*, *plc-1/PLCE1*, and *dkf-1/PRKD1-3* lowered it by roughly 80%, 75%, and 95% respectively (**Figure 6B**; adjusted *P* = 0.0094, *P* = 0.0110, and *P* = 0.0023). Therefore, exogenous muscarinic activation requires the same EGL-30/Gαq, PLC-1/PLCε, and DKF-1/PRKD cassette as infection to activate HLH-30/TFEB. Consistent with a single pathway, *dkf-1/PRKD1-3* RNAi shortened survival of infection in wild-type animals, loss of *unc-17/SLC18A3* shortened it to a similar extent, and combining the two produced no further reduction (**Figure 6C**; log-rank *P* < 0.0001 for *dkf-1*/*PRKD1-3* RNAi and *P* < 0.0001 for *unc-17*/*SLC18A3*, each against wild-type empty vector).

Together with our prior published results, these data are consistent with a bifurcated cholinergic arm of intestinal host defense (**Figure 6D**). In our working model, infection triggers acetylcholine release from neurons. Acetylcholine engages muscarinic receptors on the intestinal epithelium. In parallel to the muscarinic-WNT pathway [10], those receptors signal through PLC-1/PLCε and DKF-1/PRKD to activate HLH-30/TFEB, which drives a complementary transcriptional program that improves survival of infection without reducing pathogen burden.

## Discussion

We found that muscarinic acetylcholine signaling is required for the full HLH-30/TFEB response to infection, and that muscarinic stimulation alone is sufficient to drive HLH-30/TFEB into the nucleus in the absence of a pathogen. Cholinergic stimulation enhanced resilience to *S. aureus* infection in an *hlh-30*/*TFEB*-dependent manner. This control runs through EGL-30/Gαq, PLC-1/PLCε, and DKF-1/PRKD, the same cassette we previously placed upstream of HLH-30/TFEB during infection [20]. Thus, these results identify muscarinic acetylcholine signaling as a receptor-level input that engages this conserved pathway of HLH-30/TFEB regulation during infection. The results indicate that muscarinic activation enhances resilience to infection rather than resistance to it, consistent with the burden-independent host defense functions we described previously for both HLH-30/TFEB and the muscarinic-WNT pathway [9, 10].

Inhibition of each pathway component by RNAi or pharmacology partially reduced HLH-30/TFEB activation. The resulting defects varied in penetrance across the different tissues in the animal. This variation could mean that the requirement for this pathway in HLH-30/TFEB activation differs among tissues, with other pathways acting redundantly in some tissues and not in others. Our data cannot separate that explanation from a simpler one, that the tissues differ in how well they respond to the perturbations we used. Neurons, for example, respond poorly to RNAi delivered by feeding unless SID-1 is supplied to them [30]. For consistency, we focused our analysis on the posterior intestine. Defining where this pathway is redundant and where it is uniquely required is an important question for future work.

Although neurons are the source of acetylcholine, HLH-30/TFEB becomes active in peripheral tissues, including the intestinal epithelium. In *C. elegans*, neurons control innate immune responses in non-neuronal tissues [31, 32]. We described the same arrangement for the cholinergic control of Wnt in this animal. Infection triggers acetylcholine release from neurons. Acetylcholine acts on GAR-2/mAChR and GAR-3/mAChR in the intestinal epithelium, which signal through EGL-30/Gαq to induce Wnt expression in the same tissue, and Wnt signaling induces antimicrobial genes [10]. This arrangement is not unique to *C. elegans*, because the nervous system governs host defense in peripheral tissues across animals. In mammals, stimulation of the efferent vagus nerve reduces tumor necrosis factor production and prevents endotoxic shock [33], and inhibition of macrophage tumor necrosis factor release by acetylcholine requires the α7 nicotinic acetylcholine receptor [34]. Thus, in the absence of synapses onto the intestinal epithelium, acetylcholine works as a neuroendocrine signal that coordinates innate host defense across tissues.

The cholinergic control of HLH-30/TFEB and the cholinergic control of Wnt are therefore two acetylcholine-coordinated arms of host defense in *C. elegans*. The two arms are not redundant. Both descend from muscarinic receptors and EGL-30/Gαq, and they diverge immediately below. The Wnt arm requires EGL-8/PLCβ and PKC-1/PKCη [10], whereas HLH-30/TFEB activation requires PLC-1/PLCε and DKF-1/PRKD and proceeds even when *egl-8/PLCB1-4* and the protein kinase C genes are silenced [20]. Their outputs also differ. HLH-30/TFEB induces the antimicrobial and autophagy genes the animal needs for resilience [9], whereas the cholinergic-Wnt arm induces antimicrobial genes that protect the animal [10]. How these two arms are coordinated across the animal through time is currently unknown.

Our finding that cholinergic signaling activates HLH-30/TFEB has a counterpart in mammals, although the receptor class differs. In mammalian systems, the reported connection between acetylcholine and TFEB runs mainly through the nicotinic α7 acetylcholine receptor. Activation of the α7 nicotinic receptor drives TFEB nuclear translocation and lysosomal or autophagic programs in murine macrophages [35], in pancreatic acinar cells during pancreatitis [36], in a motor-neuron model of amyotrophic lateral sclerosis [37], and in neuronal cells challenged with α-synuclein [38]. In murine macrophages, α7 stimulation activates TFEB, but that activation is known to require the lysosomal calcium exporter MCOLN1 and the calcium-dependent phosphatase calcineurin rather than a Gαq, phospholipase C, and protein kinase D module [35]. Two further studies report nicotine changing TFEB activity by routes that were not shown to require a nicotinic receptor. In vascular smooth muscle, nicotine activates TFEB through inhibition of mTORC1 [39]. In cardiac fibroblasts, however, nicotine impedes TFEB nuclear entry by binding lactate dehydrogenase A [40]. Together, these studies establish that cholinergic stimulation can control TFEB in mammalian cells, and that the control documented so far runs through the nicotinic rather than the muscarinic receptor class. In contrast, nicotine did not activate HLH-30/TFEB in *C. elegans*. The pathway may be conserved with a switch of receptor class, the two axes may be convergent rather than orthologous, or our nicotine result may reflect the divergent pharmacology of the *C. elegans* nicotinic receptor family [41] rather than the absence of a nicotinic route.

The muscarinic side of this control is comparatively undefined in mammals. Muscarinic receptors do govern the autophagy and lysosomal machinery that TFEB controls, but the mammalian studies that show this have not tested whether TFEB is involved. Stimulation of the M2 muscarinic receptor induces autophagy in human glioblastoma cells through inhibition of mTORC1 [42], the M1 muscarinic receptor promotes autophagy in prostate cancer cells through AMPK and mTOR [43], and acetylcholine acting through muscarinic and nicotinic receptors reshapes autophagy in cardiomyocytes [44]. Muscarinic agonists and vagal stimulation temper excessive autophagy and mitophagy in cardiac tissue [45–47], an M1 receptor modulator restores autophagy signaling in an Alzheimer’s disease model [48], and G-protein-coupled receptors, including muscarinic receptors, are now recognized as regulators of autophagy [49]. Perhaps in the most direct human correlate to the positive regulation of innate immunity by acetylcholine through muscarinic receptors, acetylcholine is upregulated in psoriatic skin, where it induces muscarinic receptor M1-mediated IL-23 production by skin dendritic cells [50]. However, the source of the acetylcholine and the transcriptional effects of its upregulation were not investigated. None of these studies examined the role of TFEB or was performed in an infection setting. In this context, our investigation in *C. elegans* places a muscarinic receptor upstream of HLH-30/TFEB and connects it to a defined EGL-30/Gαq, PLC-1/PLCε, and DKF-1/PRKD pathway. Further research in mammalian systems is necessary to define the physiological conditions where these findings in *C. elegans* are relevant to human health and disease.

## Materials and Methods

### Nematode strains and maintenance

*C. elegans* strains were maintained on nematode growth medium (NGM) plates seeded with *E. coli* OP50 at 15 to 20 °C according to standard procedures [51]. The strains used were JIN2104, N2 wild type. JIN2370, *unc-17(e245)* IV. JIN2215, *unc-17(e245)* IV ; *hlh-30(tm1978)* IV ; jinIs1[*Phlh-30::hlh-30::gfp ; rol-6(su1006)*]. JIN1690, *hlh-30(tm1978)* IV ; jinIs1[*Phlh-30::hlh-30::gfp ; rol-6(su1006)*]. JIN1373, *hlh-30(tm1978)* IV.

Populations were synchronized by treating gravid adults with 10 percent bleach solution [51]. Except for *hlh-30* mutants, which do not survive the L1 starvation, eggs were held overnight in a 15 mL conical tube containing 1× M9 buffer at room temperature on a rotator. The following day L1 larvae were collected by centrifugation at 800 to 1,500 × g for 1 to 2.5 min, transferred to *E. coli* OP50-seeded NGM plates, and grown at 20 °C to the L4 stage. For *hlh-30* mutants and paired wild type controls, eggs were directly placed on *E. coli* OP50 for hatching.

### Bacterial strains and culture

*E. coli* HT115(DE3) carrying the L4440 empty vector or an individual RNAi clone was used for RNAi by feeding. *E. coli* OP50 Str^R^ was a gift of Gary Ruvkun. *E. coli* was grown overnight in LB broth (MP Biomedicals 76019-880) at 37 °C with shaking at 200 rpm.

*S. aureus* SH1000 Δ*telA::KAN* (“SH1000”, JIB0187, a gift from Lindsey Shaw) was grown overnight in tryptic soy broth (Sigma-Aldrich T8907) containing 50 µg/mL kanamycin (sulfate Sigma-Aldrich 60615) at 37 °C with shaking at 200 rpm. Ten µL of the overnight culture was spread on an 35 mm tryptic soy agar (TSA) plate (DIFCO BD 236950) containing 10 µg/mL kanamycin that had been aged for 10-15 days, and incubated for 5 to 6 h at 37 °C. Seeded plates were held at 4 °C overnight and used the following day. Plate aging and 4 °C holding temper the virulence of *S. aureus* to allow resolution of susceptibility phenotypes.

### RNAi by feeding

Single colonies of *E. coli* HT115(DE3) carrying the L4440 empty vector or an RNAi clone targeting *gar-1*, *gar-2*, *gar-3*, *egl-30*, *plc-1*, or *dkf-1* were grown for 16 h at 37 °C in LB medium containing 50 μg/mL ampicillin (sodium salt Sigma-Aldrich A9518) with constant shaking. Cultures were concentrated fivefold and 300 µL was spread on NGM plates containing 50 μg/mL ampicillin and 1 mM IPTG. For empty vector conditions the full 300 µL came from one culture. For the *gar-2* and *gar-3* double RNAi condition, 150 µL of each culture was used. For the *gar-1*, *gar-2*, and *gar-3* triple RNAi condition, 100 µL of each culture was used. Seeded plates were air dried in a fume hood and left overnight at room temperature to induce dsRNA expression. Ten gravid adult animals were then placed on each plate and allowed to lay eggs, and the progeny were grown at 15 °C for 4 days to yield L4 F1 animals for the assay.

### Infection

For infection and for the matched nonpathogenic controls, animals were placed on *S. aureus* SH1000-seeded TSA plates or on *E. coli* OP50-seeded NGM plates for 30 min at 25 °C and then processed for imaging as described below.

### Muscarinic antagonist and PRKD inhibitor treatment

Atropine (5 mM, Sigma A0257), scopolamine (5 mM, Sigma PHR1470), the selective PRKD inhibitor CRT0066101 (5 μM, Abcam ab144637), and an equal volume of ultrapure water as vehicle were each added independently to NGM plates seeded with *E. coli* OP50, and the plates were air dried. Fifty age-synchronized L4/young adult HLH-30::GFP animals were placed on each condition and incubated at 25 °C for 16 h. Fifteen to forty animals per condition were then transferred to an *S. aureus* SH1000-seeded TSA plate and infected for 30 min, washed three times in M9 buffer [51], and processed for imaging as described below. Conditions were staggered by 30 min so that every condition received the same infection time.

### Cholinergic agonist treatment

Arecoline (10 mM, Sigma 31593), oxotremorine (10 mM, Sigma O100), carbachol (10 mM, Calbiochem 212385), nicotine (hemisulfate 10 mM, Sigma-Aldrich N1019), and an equal volume of ultrapure water as vehicle were each added to NGM plates seeded with *E. coli* OP50, and the plates were air dried. After the liquid dried, young adult HLH-30::GFP animals were transferred to each plate and incubated at 25 °C for 30 min, then processed for imaging as described below. Conditions were staggered by 30 min. Animals in this assay were fed *E. coli* OP50 and were not infected.

### Muscarinic receptor knockdown during infection

Animals were reared by RNAi feeding as described above on the L4440 empty vector, the *gar-2* and *gar-3* double RNAi condition, or the *gar-1*, *gar-2*, and *gar-3* triple RNAi condition. Ten gravid adult animals were placed on each plate and the progeny were grown at 15 °C for 4 days to the L4/young adult stage. L4/young adult progeny were washed three times in M9 buffer, transferred to plates seeded with *E. coli* OP50 or *S. aureus* SH1000, and incubated at 25 °C for 30 min. Conditions were staggered by 30 min. Animals were then processed for imaging as described below.

### Arecoline treatment of animals depleted of muscarinic receptors

Animals were reared by RNAi feeding as described above on the same three conditions. L4/young adult progeny were washed three times in M9 buffer, transferred to plates containing 10 mM arecoline or vehicle, and incubated at 25 °C for 30 min. Conditions were staggered by 30 min. Animals were then processed for imaging as described below.

### Knockdown of the EGL-30/Gαq pathway

Single colonies of *E. coli* HT115(DE3) carrying the L4440 empty vector or an RNAi clone targeting *egl-30*, *plc-1*, or *dkf-1* were grown and induced as described above. Animals were reared by RNAi feeding as described above. L4/young adult progeny were washed three times in M9 buffer, transferred to plates seeded with *E. coli* OP50, plates seeded with *S. aureus* SH1000, plates containing 10 mM arecoline, or plates containing vehicle, and incubated at 25 °C for 30 min. Conditions were staggered by 30 min. Animals were then processed for imaging as described below.

### Image acquisition

Animals were anesthetized using 100 mM NaN₃ (Sigma-Aldrich S2002) on a 4 percent agarose pad on glass slides and covered with glass coverslips. Animals were imaged on a Lionheart FX automatic microscope (BioTek Instruments) using 4× or 20× objectives. Exposure and intensity settings were constant within each biological replicate. Z-stacks were automatically acquired and the best-focused plane selected for analysis. Fluorescence microscopy experiments were repeated three independent times for each treatment.

### Image analysis

Greyscale images were used for image analysis, which was performed in FIJI [52]. To measure HLH-30::GFP nuclear localization, a region of interest was drawn by hand with the segmented line tool, in the anterior end (head) or the posterior half of the intestine. The head was defined as the region from the mouth to the pharyngeal-intestinal valve. The posterior intestine was defined as the post-vulval half. RAW greyscale images were converted to 8-bit and a threshold was applied. The threshold for elimination of background fluorescence and non-intestinal nuclei was manually calibrated using positive treatment and control animals, and was held constant within a biological replicate. Regions of interest were combined in the ROI Manager using OR (Combine) and analyzed with the Analyze Particles function. Analyze Particles parameters were Size approximately 10 to 30 for the intestine, Size approximately 2 to 10 for the head, and Circularity 0.1 to 1 for both. The scorer was blinded to experimental condition.

### Bacterial burden

*S. aureus* burden was measured as colony-forming units (CFU) per animal. Five to ten animals per condition were transferred to a 1.5 mL microcentrifuge tube containing 200 µL of 25 mM levamisole (MP Biomedicals 155228) and allowed to settle, and the supernatant was removed. Animals were resuspended in 1 mL of 25 mM levamisole supplemented with 100 µg/ml streptomycin and incubated at room temperature for 30 to 60 min on a rotisserie. Animals were then rinsed three times with 1 mL of 25 mM levamisole (to arrest peristalsis) containing 1 percent Triton X-100 (to prevent animals from sticking to the plasticware, Santa Cruz sc-29112A). After the final wash, 350 µL of the same solution was added and animals were allowed to settle at room temperature. A 50 µL aliquot was taken from the top of the tube and plated to measure the extracellular bacterial load. An equal volume of carbide beads (BioSpec 11079110sc) was added to the remaining 300 µL, the suspension was vortexed at full speed for 1 min, and 30 µL was spread on triplicate TSA plates containing 10 µg/mL kanamycin by serial 10:1 dilution. CFU per animal was calculated with the following formula:

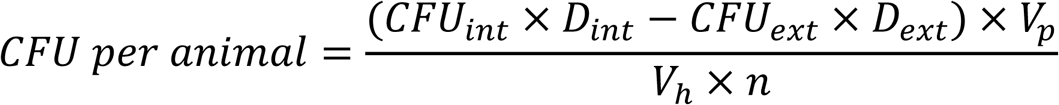

where

*CFU_int_* = internal CFU

*D_int_* = internal dilution factor

*CFU_ext_* = external CFU

*D_ext_* = external dilution factor

*V_p_* = plated volume (50 µl)

*V_h_* = homogenate volume (300 µl)

*n* = number of animals (10)

### Burden during infection of *unc-17*/*SLC18A3* animals

Thirty L4 N2 and *unc-17(e245)* animals were transferred to *S. aureus* SH1000-seeded TSA plates and held at 25 °C. Animals were harvested at 6 h and at 12 h and processed as above. At each timepoint, three tubes of ten pooled animals were processed per genotype, and the three tubes of one experiment count as one biological replicate.

### Burden after arecoline treatment

Two hundred age-synchronized L4 N2 and *hlh-30(tm1978)* animals were incubated on 60 mm NGM plates with FUdR to prevent matricidal egg hatching (100 µg/mL, Fisher Scientific 50-488-664) and held at 25 °C for 24 h. The following day, arecoline (Sigma 31593) was added to a final concentration of 5 mM (or an equal volume of vehicle) on fresh 60 mm *E. coli* OP50-seeded NGM plates and air dried. Approximately 30 animals of each genotype were transferred to these plates and held at 25 °C for 16 h, then used for infection as above.

### Survival during infection

#### Arecoline treatment

N2 and *hlh-30(tm1978)*/*TFEB* animals were synchronized and grown to the L4 stage as described above. L4 animals were transferred to fresh 60 mm *E. coli* OP50-seeded NGM plates on which 200 µL of FUdR (100 mg/mL, Fisher Scientific 50-488-664) had been spread and air dried, and were held at 25 °C. The following day, arecoline (Sigma 31593) was added to a final concentration of 1 mM (or an equal volume of vehicle) on fresh 60 mm *E. coli* OP50-seeded NGM plates and air dried. Approximately 100 FUdR-treated young adult animals of each genotype were transferred to these plates and held at 25 °C for 16 h. Animals were then transferred to 35 mm *S. aureus* SH1000-seeded infection plates and held at 25 °C. Deaths and censored events were recorded twice daily. An animal was scored as dead when it failed to move after a soft touch with the pick. Animals that desiccated on the walls of the plate or died of internal hatching were censored. Counts for the three trials are given in S1 Data.

#### Epistasis with *dkf-1*/*PRKD*

Single colonies of *E. coli* HT115(DE3) carrying the L4440 empty vector or the *dkf-1* RNAi clone were grown and induced as described above, and 300 µL was spread per plate. Adult N2 and *unc-17(e245)* animals were transferred to these plates to lay eggs, and the progeny were grown at 15 °C for 4 days to the L4 stage. FUdR (200 µL of 100 mg/mL) was then added to the plates, which were air dried and held at 25 °C for 24 h. Animals were transferred to 35 mm *S. aureus* SH1000-seeded infection plates in triplicate and held at 20 °C. Deaths and censored events were recorded twice daily. An animal was scored as dead when it failed to move after a soft touch with the pick. Animals that desiccated on the walls of the plate or died of internal hatching were censored.

### RT-qPCR

Following infection with *S. aureus* at 25°C for 8 h, animals from all conditions were washed three to four times with water and lysed in 1 mL of TRIzol reagent (Invitrogen) as previously described [10, 53]. Samples were snap-frozen in liquid nitrogen and stored at −80°C. RNA was extracted using 1-bromo-3-chloropropane (MRC) and purified by isopropanol–ethanol precipitation. Total RNA was then treated with DNase (Bio-Rad), and 250 ng was used for cDNA synthesis with the iScript cDNA Synthesis Kit (Bio-Rad). The resulting cDNA was diluted 1:5 with RNase-free water, and 2 µL of diluted cDNA was used for each RT-qPCR reaction. RT-qPCR was performed using SYBR Green Supermix (Bio-Rad) on a ViiA 7 Real-Time PCR System (Applied Biosystems). Relative gene expression was calculated using the Pfaffl method [54], with *snb-1* as the reference gene. Primer sequences are given in S1 Data.

### Statistics

Statistical analyses were performed in GraphPad Prism 10 and OASIS2 [55] and presented in S1 Data. For the fluorescence analyses, data are presented as mean ± SD, a two-sample two-sided *t* test for two groups, one-way ANOVA followed by Šídák’s or Dunnett’s multiple comparisons tests for more than two groups. Log-rank test for survival curves. For the bacterial burden experiments, each biological replicate is the mean of the log10 of its technical tubes, and the two genotypes were compared across the three replicate experiments by a paired two-tailed *t* test on those log values. * *P* ≤ 0.05, ** *P* ≤ 0.01, *** *P* ≤ 0.001, **** *P* ≤ 0.0001. ns, not significant.

## Author contributions

Swarupa Mallick: Methodology, Investigation, Visualization, Writing – review & editing. Xavier Gonzalez: Methodology, Investigation, Resources, Visualization, Writing – review & editing. Khursheed Wani: Conceptualization, Methodology, Visualization, Investigation, Writing – review & editing. Sid Ahmed Labed: Conceptualization, Investigation, Writing – review & editing. Javier E. Irazoqui: Conceptualization, Formal analysis, Funding acquisition, Methodology, Supervision, Visualization, Writing – original draft, Writing – review & editing.

## Data availability

Numerical values underlying the graphs in Figs 1 to 6 are provided in S1 Data. *C. elegans* strains and plasmids generated in this study are available from the corresponding author on request.

## Acknowledgments

We thank Lindsey Shaw for *S. aureus* SH1000 Δ*telA*::KAN (JIB0187), and Gary Ruvkun for *E. coli* OP50 Str^R^. Some strains were provided by the *Caenorhabditis* Genetics Center (CGC), which is funded by NIH Office of Research Infrastructure Programs (P40 OD010440). We thank the National BioResource Project, Japan, for *hlh-30*(tm1978).

## Supporting information

**S1 Data. Supporting numerical values.** Numerical values underlying the 18 graphed panels of Figs 1 to 6, one sheet per panel, each giving the tabulated replicate values, the summary statistics, and the statistical test reported in the corresponding figure legend. Sheets Fig5B_AllTrials and Fig6C_AllTrials also give the survival data and the log-rank statistics for every independent trial. Sheet RT-qPCR Primers gives the primer sequences used for RT-qPCR. Sheet Index lists the contents. (XLSX)

